# Genomic language models improve cross-species gene expression prediction and accurately capture regulatory variant effects in Brachypodium mutant lines

**DOI:** 10.64898/2026.02.27.708524

**Authors:** Behrooz Vahedi Torghabeh, Camous Moslemi, Julie D. Jensen, Stephan Hentrup, Tianyi Li, Xiaqing Yu, Hai Wang, Torben Asp, Guillaume P Ramstein

## Abstract

Predicting gene expression from cis-regulatory DNA sequences at the promoter and terminator regions is a central challenge in plant genomics. This capability is also a prerequisite for assessing the effects of regulatory mutations on gene expression. Here, we developed deep learning sequence-to-expression (S2E) models that leverage context-aware sequence embeddings from the PlantCaduceus genomic language model instead of one-hot encoding of sequences, to predict gene expression across 17 plant species. To further improve predictions, we integrated chromatin accessibility data as auxiliary regulatory features.

First, we evaluated our models to predict gene expression on unseen gene families via cross-validation, demonstrating our model’s prediction accuracy across all species outperforms PhytoExpr, the current state-of-the-art (SOTA) S2E model in plants (Pearson R=0.82 vs. R=0.74). We then validated variant effect predictions using an experimental dataset across 796 *Brachypodium* mutant lines, specifically designed to test predictions at single-base resolution. Our models outperformed SOTA S2E models in predicting between-gene expression differences (regression coefficient β=0.78 vs. β=0.57). Remarkably, they also accurately predicted the effects of single-nucleotide mutations on within-gene expression, while SOTA S2E models showed only weak associations (regression coefficient β=0.38 vs. β=0.08). Our results demonstrated the value of context-aware DNA sequence embeddings for predicting regulatory variant effects in plants. They also reveal a persistent accuracy gap in S2E models when moving from between-gene to allelic variation, a challenge that needs to be addressed in future S2E studies.

## 1. Introduction

A deep understanding of the mechanisms that control gene expression is an important subject in modern plant biology and a key goal for crop improvement and agricultural biotechnology [1]. The ability to predict gene expression levels from DNA sequence alone would represent a major improvement and achievement in that direction.

Gene expression regulation is mainly encoded in the non-coding DNA sequence, specifically within *cis*-regulatory elements (CREs) [2]. These CREs, such as promoters, enhancers, and silencers, are often located in the regions surrounding the transcription start site (TSS) and the transcription termination site (TTS) [3]. The divergence of CREs is a common cause of evolutionary change, i.e. mutations that affect gene expression (quantitatively or qualitatively) contribute to phenotypic diversity within and cross species. Cis-regulatory variants are a primary source of phenotypic diversity, yet the cis-regulatory code—the sequence rules governing non-coding gene regulation—remains difficult to decode, especially for non-coding variation [4], due to the complex motif grammar including spacing, orientation, and higher-order interactions that challenge predictive models [5].

Unlike traditional methods that rely on hand-engineered features (like known TF binding motifs or sequence k-mers), deep learning models can learn relevant patterns *de novo* directly from DNA sequences [6]. Recent reviews have highlighted the success of various deep learning architectures, particularly convolutional neural networks (CNNs), for DNA sequence to gene expression (S2E) modeling [6], [7]. CNNs are well-suited for this task because their convolutional filters (kernels) are capable of detecting short, spatially conserved patterns that are similar to binding motifs, and subsequent layers can learn hierarchical combinations of these features. Studies in humans have leveraged CNNs, for instance, to predict mRNA abundance directly from genomic sequences [8], to model cell-type-specific epigenetic and transcriptional profiles from sequences [9], to predict cross-species (human and mouse) regulatory sequence activity [10], and to predict gene expression using genotypic variation and functional annotations [11]. In plants, CNNs have already been used to predict gene expression quantified by transcript per million (TPM) values [1], [12], [13].

Two recent examples of S2E models in plants are PhytoExpr [1] and DeepCBA [14]. PhytoExpr predicts mRNA abundance and species from cis-regulatory DNA sequences across 17 plant species, and DeepCBA predicts gene expression in maize based on DNA sequences and chromatin interaction. Both of these models, similar to many other deep learning models, represent DNA sequences using one-hot encoding, where nucleotides A, C, G, and T are converted into binary vectors. One-hot encoding is fairly fast and straightforward to implement, can be done without requiring any specific model, and is not limited by computational costs. Yet, one-hot encoding has inherent limitations: it treats each nucleotide as an independent entity, failing to capture biochemical properties or evolutionary context. One-hot encoding does not account for the sequential orders of nucleotides and cannot measure the distance between related motifs [15]. As a result, one-hot encoding cannot represent longer-range dependencies within regulatory sequences, limiting its ability to capture relationships beyond strictly local nucleotide patterns [16].

In recent years, genomic language models (gLMs) have emerged as powerful tools that can generate sequence embeddings which can be alternatives to one-hot encodings [17], [18], [19], [20]. These types of models are trained on vast collections of datasets of unlabeled genomic sequences to learn the fundamental patterns and “language” of DNA in a self-supervised approach (without any sequence annotations). For example, the PlantCaduceus model was pre-trained on a diverse dataset of 16 angiosperm genomes and can generate context-aware sequence embeddings for any given DNA sequence [17]. In this study, we hypothesize that these pre-trained embeddings capture higher-order syntactic and semantic information from DNA sequences, providing a richer feature representation for downstream prediction tasks than one-hot encoding.

Gene expression prediction is further complicated by chromatin state, which modulates CRE accessibility [21], [22]. Regulatory features have a mutual relationship, as DNA sequences determine where TFs can bind to DNA for regulating gene expression, but TFs can also change the accessibility of chromatin, and transcription itself can also alter the chromatin state [6]. In human, incorporating information about chromatin accessibility has proved useful to predict gene expression [23], [24], [25]. In plants, even though many models have been proposed to predict gene expression, few of them have taken chromatin accessibility into account. DeepCBA used maize chromatin interaction data to train a deep learning model for predicting gene expression and achieved a higher accuracy in expression classification and expression value prediction compared to existing models [14]. Therefore, we hypothesize that augmenting sequence information with chromatin accessibility information generated here using the cross-species model a2z will significantly improve prediction accuracy in gene expression [26].

In plants, different approaches have been used to validate recent S2E models. For instance, [1] and [12] have validated their models by showing that their models are able to learn biologically meaningful regulatory motifs from DNA sequences. Authors of [1] have also used a statistical approach for model validation and have shown that there is a significant association between model predictions and the effects of expression quantitative trait loci (eQTLs) in natural populations. Some studies such as [1], [27], [28] have leveraged experimental validation *in vitro* and quantified the functional impact of designed or altered sequences using transient expression assays in plant protoplasts. To our knowledge, S2E models have not yet been validated for their ability to predict the effects of single point mutations in whole plants (*in planta*). In this study, we address this critical gap by using a novel mutant population to test the ability of S2E models to predict the effect of point mutations [29].

PhytoExpr established a strong baseline for predicting gene expression across plant species from cis-regulatory DNA sequences around promoter and terminator regions using one-hot encoding of TSS and TTS sequences [1]. Building on that, we developed new S2E models that replace one-hot encoding with context-aware sequence embeddings from the PlantCaduceus gLM while also incorporating sequence embeddings and chromatin accessibility predictions from the a2z model for the same TSS and TTS regions. We trained our models on approximately 0.6 million genes across 17 plant species (same dataset as [1]) by using dedicated CNN architectures to predict median TPM values (Fig. 1). We then perform a two-tiered evaluation. First, we assessed cross-species predictive performance via cross-validation (CV) evaluation against PhytoExpr. Second, and more importantly, we conducted rigorous *in planta* validation using a novel, purpose-built population of 796 mutagenized and selfed *Brachypodium* lines (SIEVE). This population was uniquely designed to test variant effect predictions at single-base resolution. Our work thus provides: (i) a framework for leveraging gLMs and integrative regulatory features for expression prediction, and (ii) a benchmark for the challenging task of predicting single-nucleotide variant effects in plants.

**Figure 1:**
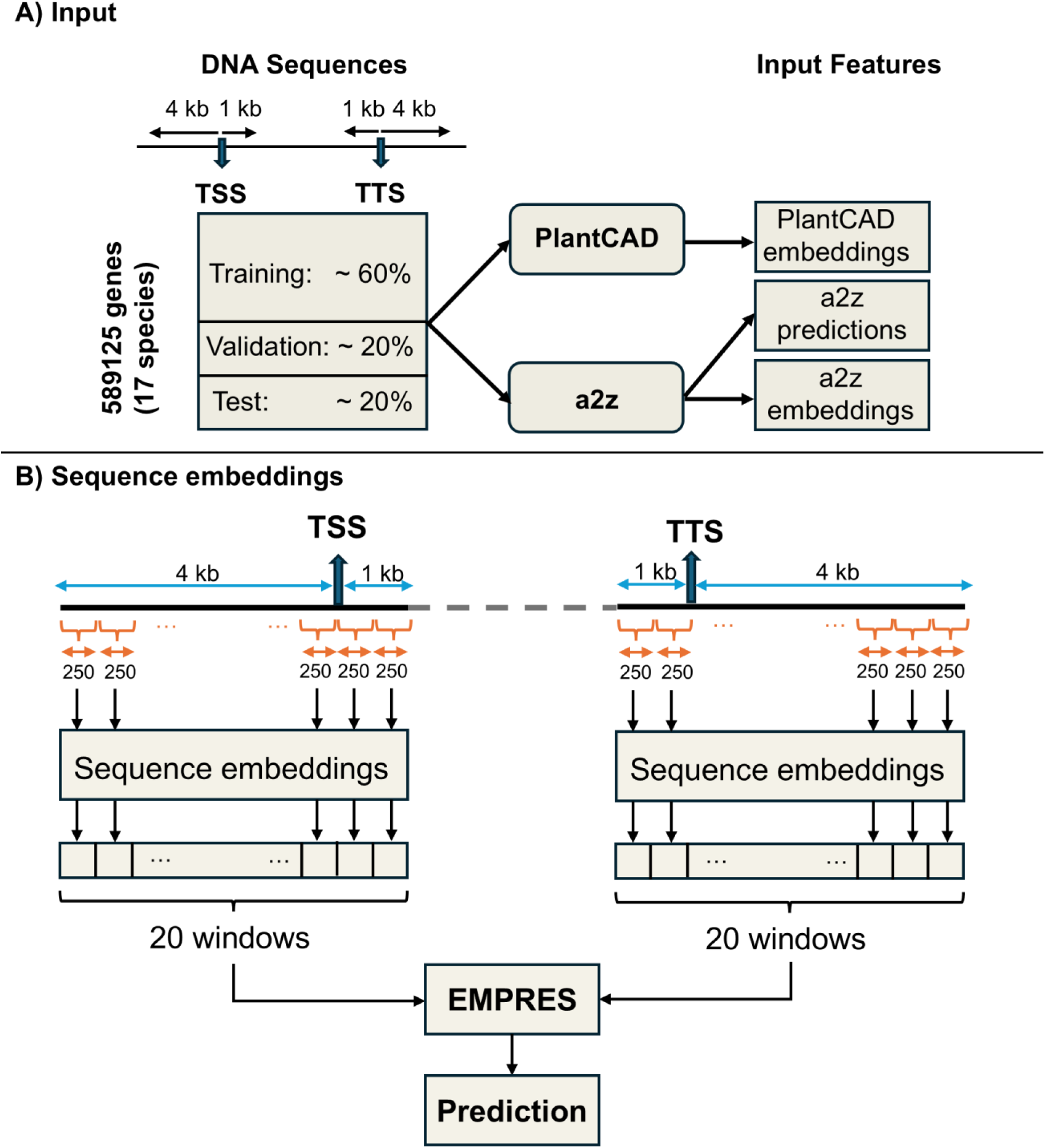
**a) Overview of the input data and feature generation pipeline**. For each gene across 17 angiosperm species (589,125 genes), two 5000 bps (5 kb) regulatory DNA sequences were extracted: one centered on the transcription start site (TSS; 4 kb upstream, 1 kb downstream) and one centered on the transcription termination site (TTS; 1 kb upstream, 4 kb downstream). Genes were split into training (∼60%), validation (∼20%), and test (∼20%) sets based on the same gene-family-aware setup as PhytoExpr. Regulatory sequences were passed to pretrained PlantCaduceus and a2z models to generate sequence embeddings and chromatin accessibility-related features, which served as inputs to the EMPRES models. **b) Generating regulatory sequence embeddings.** Each TSS- and TTS-centered sequence was divided into 20 overlapping windows. For each window, a 250 bp core region was fed to PlantCaduceus and a2z, with flanking regions providing contextual buffering. PlantCaduceus embeddings were averaged across the core sequence to produce one vector per window (20 × 384 per region), whereas a2z embeddings were extracted directly from the penultimate layer (20 × 925 per region). Embeddings from all windows spanning the TSS and TTS regions were concatenated and used by EMPRES to predict gene expression. PlantCAD: PlantCaduceus genomic language model

**Figure 2:**
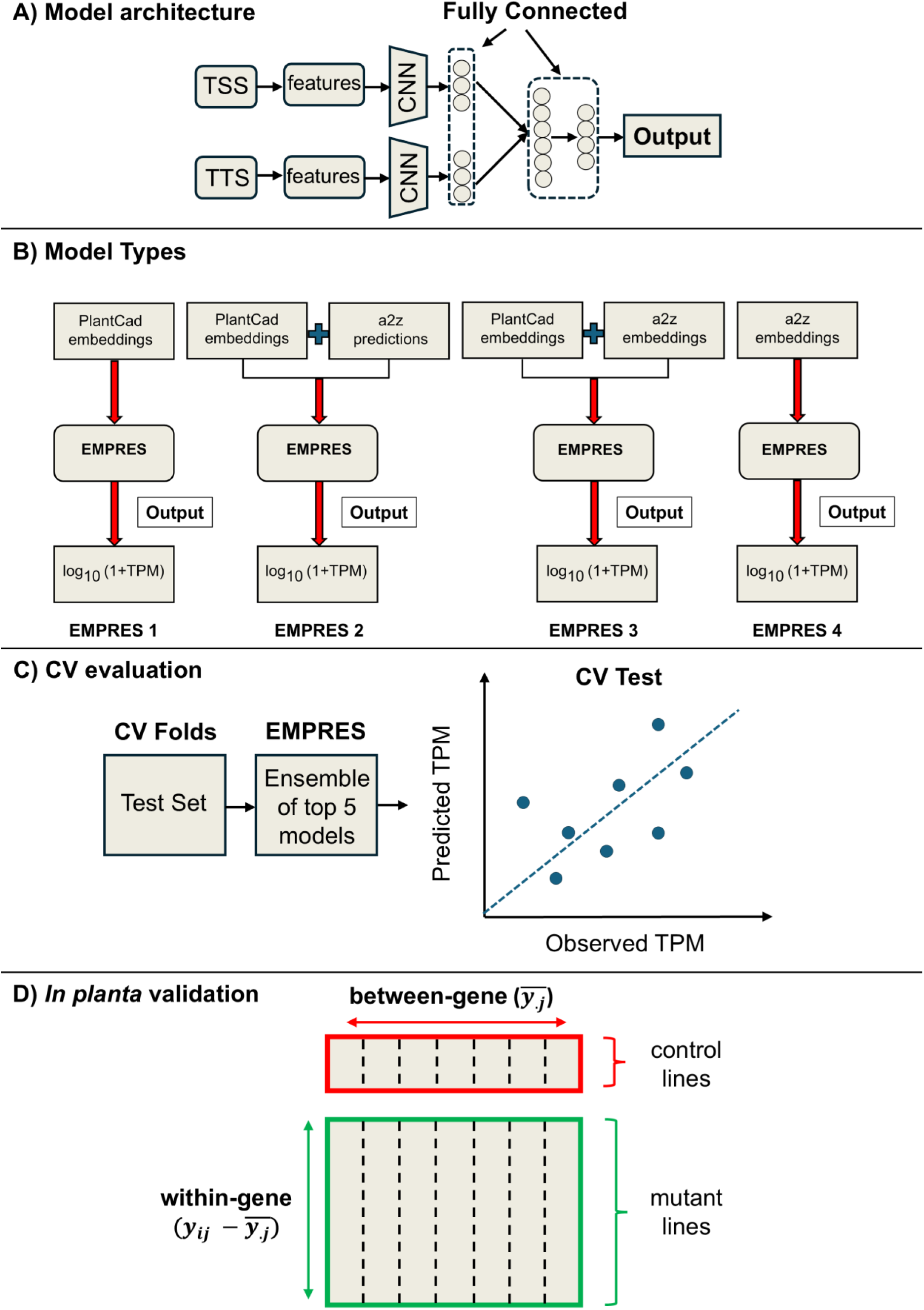
EMPRES model architecture, model variants, and evaluation strategy. **a)** Schematic of the EMPRES model architecture. Regulatory features derived from TSS and TTS sequences are processed in two parallel branches using identical 1D convolutional neural networks and fully connected (dense) layers. Outputs from the TSS and TTS branches are concatenated and passed through fully connected layers to predict gene expression. **b)** Overview of the four EMPRES model variants and their corresponding input features. EMPRES 1 uses PlantCaduceus embeddings; EMPRES 2 combines PlantCaduceus embeddings with a2z chromatin accessibility predictions; EMPRES 3 combines PlantCaduceus embeddings with a2z embeddings; and EMPRES 4 uses a2z embeddings alone. All models predict log_10_ (1+TPM). **c)** Cross-validation (CV) evaluation strategy. Models were trained using five-fold CV with predefined gene family-based splits. For each fold, the top five models (selected by validation loss) were ensembled to generate predictions on the held-out test set, and predictive performance was assessed by comparing predicted and observed TPM values. **d)** *In planta* validation framework using the SIEVE mutant population. Between-gene variation was evaluated using mean expression across control lines for each gene, whereas within-gene (allelic) variation was assessed by deviations of mutant expression from the corresponding gene-specific control mean.

## 2. Materials and methods

### 2.1 Genomic and transcriptomic data across angiosperms

#### 2.1.1 Source data and species coverage

The baseline dataset in this study is the published data for PhytoExpr (data.csv) [1] which contains 10,000 base pairs (bp) DNA sequence centered around both the TSS and TTS of genes for 17 angiosperm species covering an evolutionary timescale of 150 Mys. The dataset also denotes species, gene ID, gene family, and expression values (TPM) for each gene. Each row in the source dataset with valid TSS and TTS sequences were processed by extracting two 5,000 bp sequences, one 4,000 bp upstream and 1,000 bp downstream of TSS, and the other 1,000 bp upstream and 4,000 bp downstream of TTS.

### 2.2 Sequence Representations

#### 2.2.1 Generating regulatory DNA sequence embeddings

Each TSS and TTS sequence was divided into 20 overlapping windows with varied size depending on the models for downstream tasks. A window size of 512 was used for PlantCaduceus, while a2z has a window size of 600. Each window consists of a core sequence of 250 bp, placed in the middle, with the flanking regions corresponding to the remainder (2×131 for PlantCaduceus and 2×175 for a2z) and serving as buffer for smoother transitions between each of the 20 successive windows. Sequence embeddings from both PlantCaduceus and a2z were extracted from the penultimate neural network layer for each of the 20 partitions and saved along with species, gene ID, gene family and TPM values. Python v3.11.7 was used for RNA seq analysis, sequence extraction and generating sequence embeddings.

#### 2.2.2 PlantCaduceus Embeddings

Embeddings were extracted from the “PlantCaduceus_l20” model which features an input size of 512 bp and 384 embeddings for each input position [17]. For each of the 20 TSS and TTS input windows, only the embeddings for the 250bp core sequence were used (250×384). The embeddings for each window were then pooled into one average embedding, resulting in a final embedding set of size 20×384 for the TSS sequence, and 20×384 for the TTS sequence.

#### 2.2.3 a2z Embeddings

The a2z model is a deep learning model trained on 12 angiosperm species, which can predict chromatin accessibility and DNA methylation from DNA sequences across plant species [26]. Embeddings were extracted from the “model-accessibility-full.h5” a2z model with an input size of 600 base pairs, a penultimate layer size of 925 and a predicted output size of 1 [26]. Unlike PlantCaduceus, a2z does not generate one embedding for each base pair in the input. Therefore the 925 embeddings for each 20 TSS and TTS windows were used to form the final embedding set of size 20×925 for the TSS sequence and 20×925 for the TTS sequence. Additionally, the chromatin accessibility prediction from the final layer window was saved separately for each window (20×1).

#### 2.2.4 Data Preprocessing: Cross-validation splits and standardizing

First, gene families and their corresponding genes were divided into the same 5 disjoint groups as in [1] for 5-fold CV. Then, the training data (PlantCaduceus embeddings and a2z predictions and embeddings) were integrated into one file for each species. Sequence features consist of two tensors (one tensor for TSS and one for TTS) of shape (N, C, L). For each tensor, N corresponds to 589,125 genes across all species, C is the number of channels (features) at each position, equal to 384 for PlantCaduceus embeddings, 1 for a2z predictions, and 925 for a2z embeddings, and L is the number of positions (locations), equal to 20 for our proposed model and 5,000 for PhytoExpr.

The CV fold pairs were set as (1, 2), (2, 3), (3, 4), (4, 5), and (5, 1), where the first and second elements in each pair denote the validation and test group numbers, respectively. The remaining groups in each pair were used as the training set. These splits ensure each group of genes is left out exactly once for being used in CV tests. Based on the CV splits, standardization metrics (mean and standard deviation) were calculated from the corresponding training set for each channel at each position for each TSS and TTS sequence in each CV fold. Subsequently, features were standardized to zero mean and unit variance in the training set. Lastly, all TPM values were transformed to log_10_ (1+TPM), ranging from 0 to 4.12.

### 2.3 Prediction models

#### 2.3.1 Model types and their corresponding inputs

The training data and transformed TPM values were further used in the downstream analysis for training four types of models, performing CV tests, and generating predictions of TPM values. The models and their corresponding input data were called ‘Embedding-based Prediction of Expression from Sequence’ (EMPRES) 1, 2, 3, and 4. EMPRES 1 models were trained on PlantCaduceus embeddings, EMPRES 2 models were trained on PlantCaduceus embeddings and a2z predictions, EMPRES 3 models were trained on PlantCaduceus embeddings and a2z embeddings, and EMPRES 4 models were only trained on a2z embeddings. The input data dimensions for EMPRES 1, 2, 3, and 4 models were (N, 384, 20), (N, 384+1, 20), (N, 384+925, 20), and (N, 925, 20) respectively.

#### 2.3.2 Model architecture

In order to train a deep learning model on regulatory sequences around the promoter and terminator regions to predict gene expression, we developed a custom dual branch 1D convolutional neural network. This model is designed to take data for TSS and TTS in two separate branches, referred to as TSS branch and TTS branch. Each branch in our custom CNN has the same architecture and consists of 3 to 5 1D convolutional layers followed by 3 to 5 fully connected layers, selected through hyperparameter optimization (Table S1). Then, the outputs of TSS and TTS branches (a 1D vector) were concatenated into one unified vector and passed through additional fully connected layers for further processing, with the number of post-merge layers ranging from 2 to 4 (Table S1). The number of input channels in our CNN varies depending on the type of models and their corresponding input data. Each model was trained using PyTorch 2.5.1 [30] with CUDA 12.4 and cuDNN 9.1.0

#### 2.3.3 Hyperparameter optimization using Optuna

Before training the EMPRES models, the hyperparameter values were tuned on each data type by using the “Optuna” hyperparameter optimization software framework [31]. The hyperparameters were sampled by Optuna 4.2.1 using the TPE sampler with default parameter values and with fixed random state (for reproducibility). Hyperparameters were continuous for the learning rate and dropout, and categorical otherwise (Table S1). For each four types of models in this study, hyperparameters were optimized independently, allowing Optuna to select the best hyperparameter values based on the input data. For all models, batch size and activation functions were set to 256 and ReLU, respectively. In both branches, batch normalization was applied after the activation function in each convolutional block, and no batch normalization was applied to any of the fully connected layers.

#### 2.3.4 Model training

For each model type, 200 models were trained with different sets of hyperparameters, and the performance of each model in training and validation was measured by mean square error (MSE) loss. The “AdamW” optimizer was used for minimizing the prediction loss in training, which separates the weight decay from the gradient update [32]. During training, early stopping was used with patience of 10 epochs and minimum validation loss improvement of 0.01, with the maximum number of epochs set to 50 epochs. Each model was saved at the ‘best epoch’ where the validation loss at that epoch was minimized. After all the 200 models finished training, the trained models were saved at the best epoch, i.e. the epoch at which validation MSE loss was minimum. This pipeline was repeated for each of the five CV folds and model types. All model training was performed using a single NVIDIA L40S GPU on the GHPC cluster at the Center for Quantitative Genetics and Genomics, Aarhus University.

#### 2.3.5 Cross-validation tests and generating predictions on reference data

After model training, the top five models in each CV fold were fitted to the standardized test set data to generate predicted TPM values for each gene. Then, the top five models were ensembled to generate an average predicted TPM for each gene. Subsequently, in each CV fold the MSE test loss for each model as well as the ensemble (average) of the top five models was calculated. Finally, an overall test loss across all five CV folds was calculated for each model type to measure the accuracy of predicted TPM values. In addition, the Pearson correlation coefficients between observed TPM values and predicted TPM values from each of the top five model predictions and the ensemble of the top five were calculated for each model type to measure the effectiveness of ensembling (Supplementary Figure S3). Finally, the Pearson correlation coefficient between the average predictions across all five CV folds and the observed TPM values were calculated over all four model types. For comparison, PhytoExpr models were evaluated using the same CV splits and test sets, including the transformer-based PhytoExpr B, which predicts median TPM only, and the transformer-based PhytoExpr C, which predicts median TPM and species [1].

### 2.4 Experimental validation of model predictions in a *Brachypodium distachyon* mutant population

#### 2.4.1 Plant material

A mutant population in the model grass *Brachypodium distachyon* was developed for ‘selection of mutations by *in silico* and experimental variant effects’ (SIEVE). The SIEVE population was designed to validate predicted variant effects *in planta* at single-base resolution, as described previously (Moslemi *et al.* 2026). Briefly, seeds from *Brachypodium distachyon* accession Bd21-3 were treated with 0 mM (controls) or 7 mM (mutants) of the mutagen sodium azide at the M_1_ generation (first generation after seed mutagenesis). Then, plants were grown in the greenhouse (20h/4h light/dark, 21°C/18°C target temperature, at Aarhus University Flakkebjerg) and allowed to self-fertilize naturally across 5 generations, from M_1_ to M_5_ (Fig. 3a). At the M_5_ generation, juvenile leaf tissue was collected (about 150 mg of leaf tissue, 5 weeks after sowing), for subsequent DNA extraction and whole-genome sequencing by DNBSEQ (paired-end sequencing, 150-bp reads) at BGI, Hong-Kong. Sequencing reads (fastq files) were aligned to the *Brachypodium distachyon* Bd21-3 v1.2 assembly from Phytozome (BdistachyonBd21_3_537_v1.0.fa) by BWA-MEM [33]. SNP variants were then called by GATK [34]. Variant sites were filtered based on GATK’s standard filters (QD < 2, QUAL < 30, SOR > 3, FS > 60, MQ < 40, MQRankSum < -12.5, ReadPosRankSum < -8). The resulting VCF file contained genotype calls for 857 lines (829 mutants, 28 controls) and 581,803 variants.

**Figure 3:**
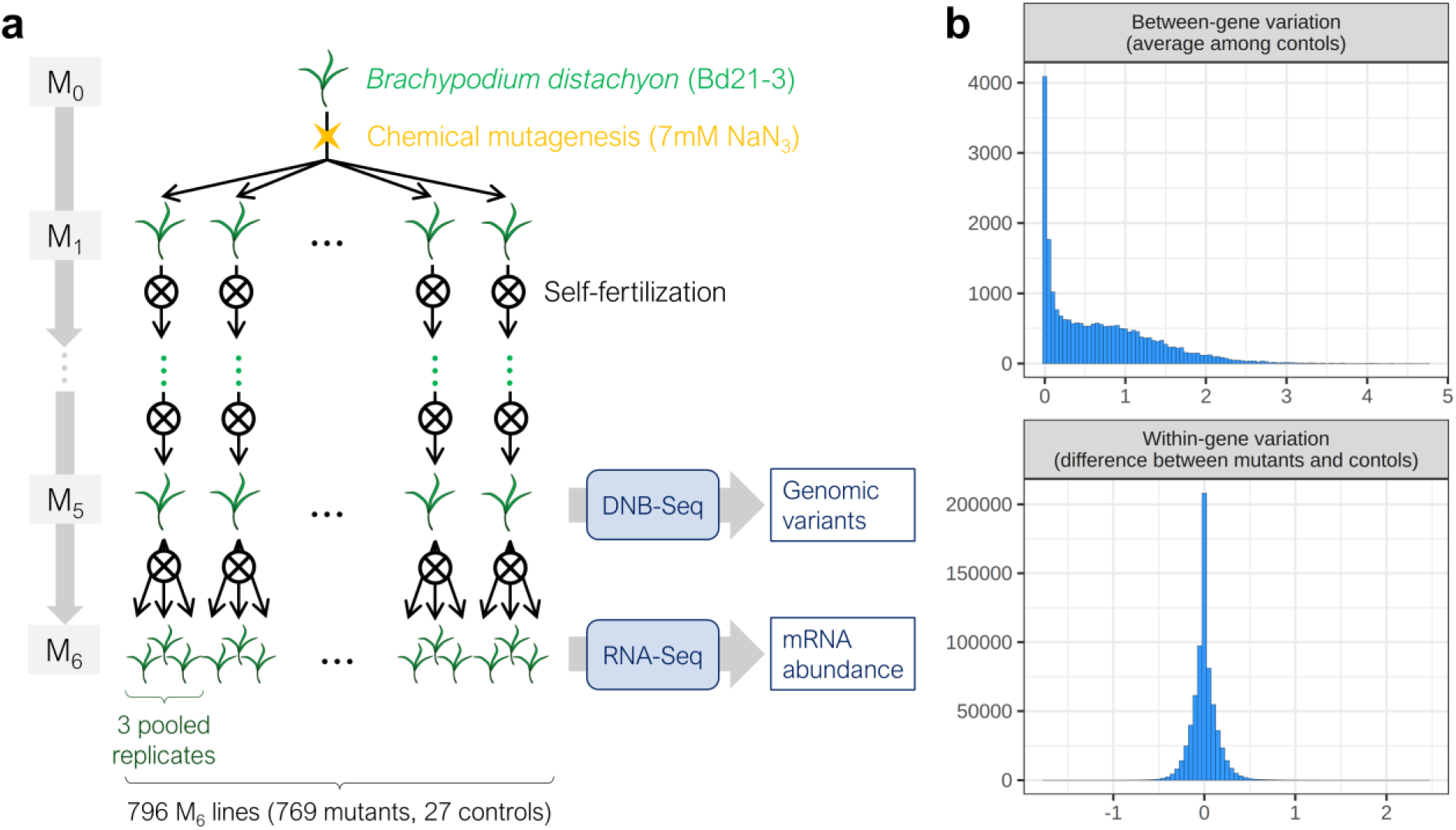
SIEVE population for experimental validation in planta. **(a)** Generation of the SIEVE population in *Brachypodium distachyon* accession Bd21-3. Seeds were treated with sodium azide (7 mM; controls: 0 mM) at the M₁ generation and self-fertilized for five generations (M₁ - M₅). Whole-genome sequencing of M₅ plants identified single-nucleotide variants, and RNA sequencing quantified gene expression (mRNA abundance) in leaf tissues at the three-leaf stage, across 796 lines (769 mutants, 27 controls). **(b)** Measurement of observed gene expression in the SIEVE population: between-gene differences (average among control lines, for each gene) and within-gene differences (difference between measured expression and average among controls, for each mutant line and gene). Gene expression was quantified as transcripts per million (TPM). SIEVE: Selection of mutations by In Silico and Experimental Variant Effects.

#### 2.4.2 Observed mRNA abundance

Seeds from each M₅ line were used to produce M₆ generation seedlings under controlled growth conditions. Prior to sowing, M₆ seeds were pre-incubated for one week at 37°C and 20% relative humidity to enhance and synchronize germination [35]. Germination and seedling growth were carried out in climate-controlled chambers at Aarhus University Flakkebjerg under a 16h/8h light/dark photoperiod, 23°C/21°C day/night temperatures, and 60% relative humidity. At the three-leaf stage (approximately 16 days post-germination), the blades of all three fully expanded leaves (excluding the cotyledons) were harvested and immediately frozen in liquid nitrogen, then stored at –80 °C until further processing. Tissue sampling was performed in three independent biological replicates per line. The three replicates from each line were subsequently pooled for total RNA extraction and RNA sequencing. After excluding the insufficient plant material (less than 3 biological replicates), 796 lines (769 mutants, 27 controls) were assayed by RNA sequencing (Fig. 3a). Library preparation was performed using Poly(A) selection and paired-end sequencing (2 × 150 bp) was conducted on an Illumina NovaSeq platform at Azenta GeneWiz (Leipzig, Germany).

The quality of paired-end reads for each line in the SIEVE population was controlled by FastQC and fastp using default options. Reads were subsequently aligned using hisat2, with default options [36] and converted into bam files using samtools [37]. The number of reads were counted using the featureCounts function of subread [38] configured to count fragments instead of reads (-p), using the Bd21-3 genome annotations (-a BdistachyonBd21_3_537_v1.2.gene.gff3). The resulting aligned feature count files (*.FC) were subsequently read using a custom R script (RNAseq.R) to calculate TPM for each line and gene, for each line in the population, which was used in downstream analyses.

#### 2.4.3 Generating sequence embeddings and predictions in the SIEVE population

TSS and TTS sequences were generated for each of the 23,725 genes of each line in the SIEVE population, similarly to the input data in the model training phase. These sequences incorporated any unique variants that each plant carried in their respective genomic TTS and TSS sites. These sequences were then used to predict TPM values for each gene, either directly with PhytoExpr or through extracted PlantCaduceus and/or a2z features with our EMPRES models. For generating the predicted TPM values in SIEVE population for each gene in a given CV fold, the embeddings were first centered and scaled using the mean and standard deviation calculated from the corresponding training set (in the PhytoExpr dataset). Then, predictions were generated by the same top five models of each type that have not seen the genes in the CV fold during training. The predicted TPM values were then compared to the actual measured TPM values from RNA-seq reads in the SIEVE population.

#### 2.4.4 Between-gene expression variation in SIEVE population

To investigate the differences in expression levels between genes, we analyzed the average gene expression among control lines (Fig. 3b). Let M be the set of predictive models tested in this study, where M = {PhytoExpr B, PhytoExpr C, EMPRES 1, …, EMPRES 4}. First, we defined the average expression for line *i* and gene *j* as *y*_*ij*_ = *log*_10_(1 + *TPM*_*ij*_). Next, the predictions on SIEVE population were filtered to include only the control samples. Then, for each gene *j*, we calculated the mean observed expression value across all N_C_ control line replicates, denoted as *y̅*._*j*_, and also calculated the mean predicted expression value, denoted as 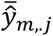. Subsequently, for each model *m* ∈ M, a separate simple linear regression model was fitted to the gene-level averages. The statistical model is specified as:

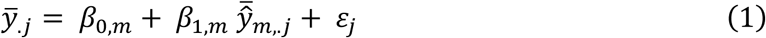

where for gene *j*, *y̅*_.*j*_ is the average expression, 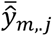 is the average predicted expression value by model *m*, *β*_0,*m*_ is the intercept, *β*_1,*m*_ is the regression coefficient for the predicted values by model *m*, and *ε*_*j*_ is the error term. Thus, *β*_1,*m*_ quantifies the prediction accuracy of model *m* for between-gene expression variation across control lines. For each fitted model, we extracted the coefficient estimate, standard error, *R*^2^, and *t*-test results (statistic and *p*-value) from the *lm* function in R.

#### 2.4.5 Within-gene expression variation in SIEVE population

To test whether our predictive models could explain allelic differences in gene expression within individual genes (i.e., deviation between mutant and control conditions), we implemented a multiple linear regression framework designed to isolate the biological signal from technical variation and latent batch effects. The analysis was performed exclusively on the mutant samples. First, to adjust for technical and non-genetic sources of variation, we determined the optimal number of principal components (PCs) to include as covariates in our models. We fitted a series of 20 null models, each regressing the observed expression change against an increasing number of PCs, as follows:

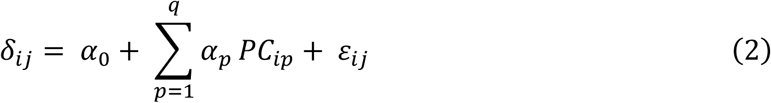

where *δ*_*ij*_ is deviation of observed gene expression (calculated as the deviation of the mutant observed expression from the gene’s average observed expression in controls: *δ*_*ij*_ = *y*_*ij*_ − *y̅*_.*j*_) and *q* represents the number of PCs included, ranging from 1 to a maximum of 20. We calculated the Bayesian Information Criterion (BIC) for each of the 20 fitted null models and selected the optimal number of PCs, denoted as *k*, that yielded the minimum BIC value.

Having determined the optimal number of covariates (*k* = 18), we then fitted a separate multiple linear regression model for each predictive model *m* ∈ M to test its explanatory power as follows:

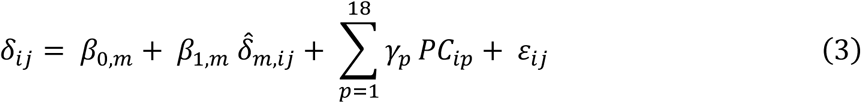

where for each gene *j* in a given mutant sample *i*, *δ*_*ij*_ is the observed change in expression, *δ*^_*m*,*ij*_ is the predicted change in expression according to model *m* (calculated as 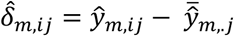), *PC*_*ip*_ represents the value of the *p*-th principal component for line *i*, *β*_0,*m*_, *β*_1,*m*_, and *γ*_*p*_ are the coefficients for the intercept, the predicted change, and the principal components, respectively, and *ε*_*ij*_ is the error term. From each final fitted model, we extracted the coefficient estimate for the predictor of interest (*β*_1,*m*_), its standard error, partial *R*² (proportion of variance explained by predicted change in expression), and *t*-test results (statistic and *p*-value). The prediction accuracy of each model for predicting the effects of allelic variation on expression is quantified by the estimated *β*_1,*m*_ regression coefficient.

## 3. Results

### 3.1 Hyperparameter optimization and the benefit of ensembling

Optuna’s parameter importance plots reveal that learning rate is the most impactful hyperparameter in optimizing the hyperparameters of our proposed models. The relative importance of learning rate in optimizing the performance (validation MSE loss) is equal to 0.73 for EMPRES 1, 0.88 for EMPRES 2, 0.90 for EMPRES 3 and 0.91 for EMPRES 4 (Figure S1). The parameter importance plots show that sampling the learning rate from the continuous interval of [10^-5^, 10^-2^] helps the model optimization, especially for EMPRES 2 and 3 models (Figure S1). Optuna Slice plots demonstrate the distribution of hyperparameter values and the corresponding validation MSE loss for each model, showing that the optimal values for learning rate for our proposed models is usually around 10^-4^ (Figure S2). Moreover, the Pearson correlation coefficients between observed and predicted TPM values from each of the top five models predictions and the ensemble of top five show that ensembling slightly increases the prediction accuracy by 0.01 compared to the top performing model for each type of our proposed models (from 0.81 to 0.82 for EMPRES 1 and 2, from 0.79 to 0.80 for EMPRES 3, from 0.68 to 0.69 for EMPRES 4). Thus, ensembling proves to be useful but to a limited extent (Figure S3).

### 3.2 EMPRES models achieve high accuracy for gene expression prediction and generalize to unseen gene families

To predict the expression from proximal regulatory sequences in promoter and terminator regions across 17 plant species, we trained supervised models consisting of two parallel CNNs with the same architectures to predict median TPM values. The results of our proposed models show that we can accurately predict median TPM values between genes across all species using regulatory sequence embeddings and open chromatin predictions as features (Figure 4a). The high correlation between the predicted and observed TPM values in left-out gene families suggests that our proposed models capture the regulatory code of gene expression using the proximal sequences in promoter and terminator regions to predict mRNA abundance (Figure 4b). Such prediction accuracy is obtained by relatively shallow deep learning models.

**Figure 4:**
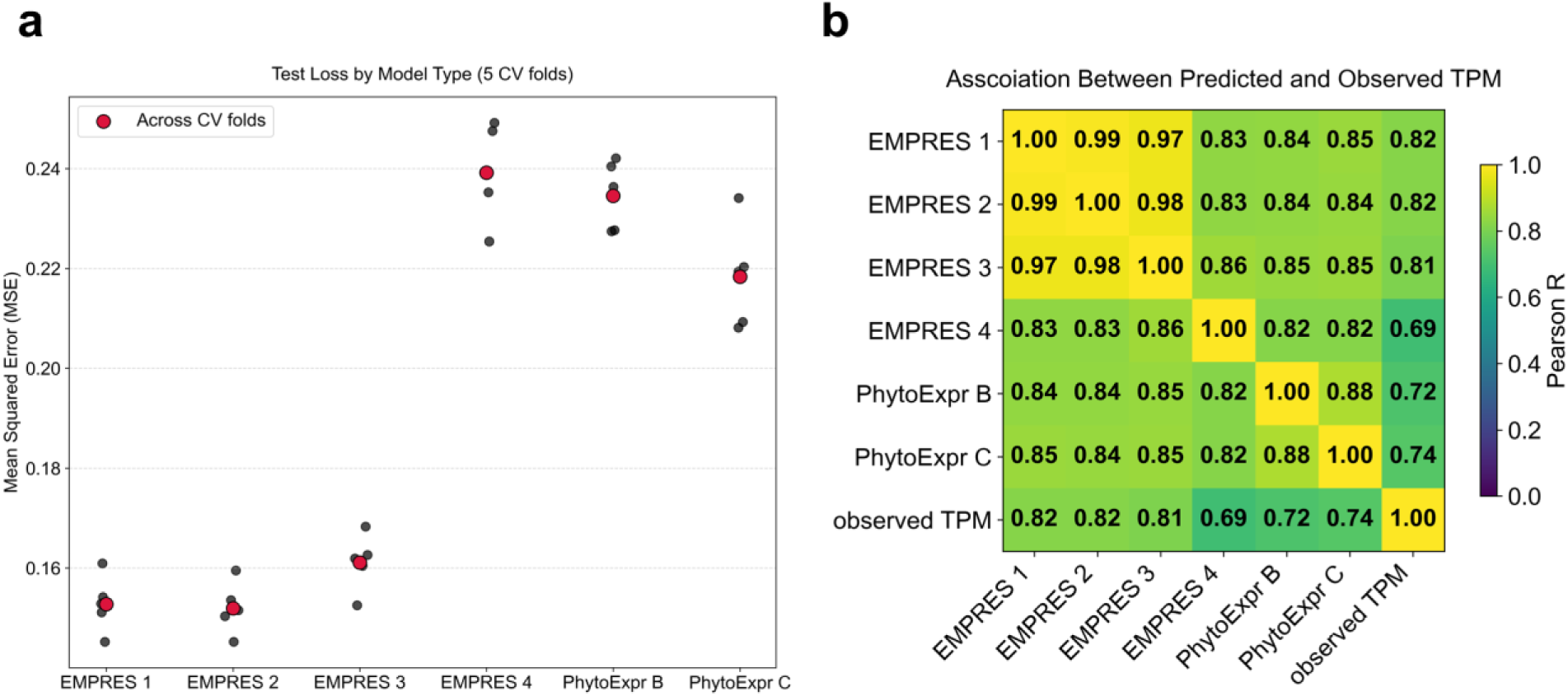
Cross-validation performance and prediction accuracy of each model. **(a)** Mean squared error (MSE) in test set across five cross-validation (CV) folds for each model. Black points denote MSE loss in each CV fold and red points denote the mean MSE loss across CV folds. **(b)** Prediction accuracy quantified by pairwise Pearson correlation coefficients (Pearson R) between model predictions and observed expression (observed TPM). EMPRES 1 uses PlantCaduceus embeddings; EMPRES 2 uses PlantCaduceus embeddings and a2z chromatin accessibility predictions; EMPRES 3 uses PlantCaduceus embeddings and a2z embeddings, EMPRES 4 uses a2z embeddings. All EMPRES models predict median TPM. PhytoExpr B and C are two PhytoExpr transformer designs, where design B predicts median TPM and design C predicts both median TPM and species. TPM: transcript per million.

The CV evaluations show that EMPRES models achieve high overall prediction accuracy across all 17 species in a gene-family-aware setting. The Pearson correlation between predicted and observed TPM values is equal to R = 0.82 for EMPRES 1 and EMPRES 2, R = 0.81 for EMPRES 3, and R = 0.69 for EMPRES 4 (Figure 4b). In comparison, the benchmark PhytoExpr transformer models achieve substantially lower accuracy (PhytoExpr B: R = 0.72, PhytoExpr C: R = 0.74), demonstrating improved generalization of EMPRES models to unseen gene families.

Among the models leveraging PlantCaduceus embeddings, EMPRES 1 and EMPRES 2 show very similar test performance and slightly outperform EMPRES 3 on the reference data (mean MSE loss = 0.15 for EMPRES 1 and 2, compared to 0.16 for EMPRES 3; Figure 4a). EMPRES 1 and EMPRES 2 therefore achieve the highest overall prediction accuracy among all evaluated models (Figure 4b). Across all species, EMPRES 1 and EMPRES 2 explain substantially more variance in gene expression (R^2^ = 0.67) than the best benchmark model (PhytoExpr C: R^2^ = 0.54).

The lower prediction accuracy of EMPRES 4 models compared to other EMPRES models is due to using only sequence embeddings generated by a2z model, which is designed to predict accessible chromatin region from DNA sequence and is not a gLM. Interestingly, the accuracy of EMPRES 4 model is still relatively close to benchmark models. This suggests that sequence determinants of chromatin accessibility are useful to predict mRNA abundance across angiosperms but are not as powerful as sequence representations from language models [6].

Across all EMPRES models, predicted expression values show a clear association with observed TPM, with the highest density of observations concentrating along the diagonal, indicating overall consistency between predictions and measurements (Figure 5a). Higher observed TPM values tend to be underpredicted, whereas low observed TPM values tend to be overpredicted, and differences in prediction density across models reflect variation in predictive accuracy. To further quantify model performance across different expression states, we evaluated prediction error separately for expressed (observed TPM ≠ 0) and unexpressed (observed TPM = 0) genes using MSE (Table 1). For both subsets, EMPRES models leveraging PlantCaduceus embeddings (EMPRES 1–3) achieve lower MSE than the benchmark PhytoExpr models, with EMPRES 1 and EMPRES 2 consistently showing the best performance, whereas EMPRES 4 and both PhytoExpr models show higher error. These results indicate that the performance advantage of EMPRES models is maintained across genes with different expression states.

**Figure 5:**
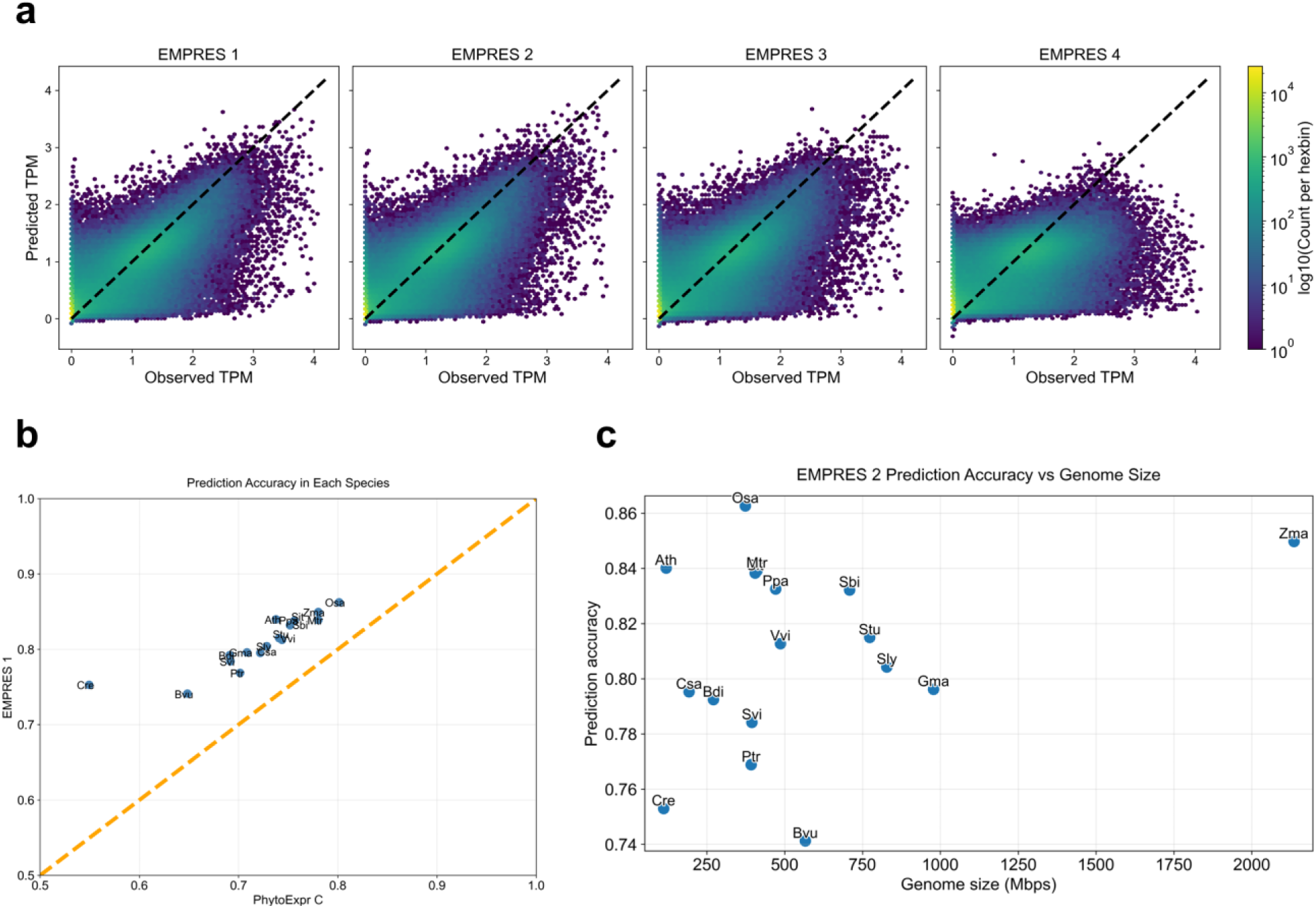
Model prediction performance across species and genomic contexts. **(a)** Hexbin density plots showing the distribution of observed versus predicted gene expression (TPM) across all species for the four proposed models (EMPRES 1 to EMPRES 4). Color intensity indicates the density of observations, ranging from 1 to 10000. **(b)** Species-specific prediction accuracy for EMPRES 2 and PhytoExpr C across the 17 plant species included in this study, measured as the Pearson correlation coefficient (Pearson *R*) between predicted and observed TPM values. **(c)** Relationship between prediction accuracy of the best-performing model (EMPRES 2) and genome size across species. Each point represents one species. Genome size is measured in million base pairs (Mbps). TPM: transcript per million.

**Table 1:** Prediction error (MSE) of EMPRES models and benchmark PhytoExpr models evaluated separately on expressed (observed TPM ≠ 0) and unexpressed (observed TPM = 0) genes. Lower MSE indicates better predictive accuracy. MSE: mean squared error.

| Expression state | EMPRES 1 | EMPRES 2 | EMPRES 3 | EMPRES 4 | PhytoExpr B | PhytoExpr C |
| --- | --- | --- | --- | --- | --- | --- |
| Expressed | 0.187 | 0.186 | 0.196 | 0.284 | 0.281 | 0.267 |
| Unexpressed | 0.074 | 0.073 | 0.080 | 0.135 | 0.126 | 0.105 |

All our models converge rapidly, with optimal validation loss typically reached within fewer than 10 epochs (Figure S4). Early stopping (patience = 10 epochs, minimum validation-loss improvement = 0.01) prevents overfitting and reduces unnecessary training. A separate validation set comprising ∼20% of all genes ensures model optimization over a sufficiently large and diverse set of gene families. Training times per CV fold were ∼34 hours for EMPRES 1 (≈171 hours total), 35.5 hours for EMPRES 2 (≈177 hours total), 57.5 hours for EMPRES 3 (≈288 hours total), and 38 hours for EMPRES 4 (≈190 hours total), using a single NVIDIA L40S GPU. Overall, this training strategy enabled efficient optimization and remained compatible with single-GPU computational constraints.

### 3.3 EMPRES models predict gene expression within species independently from genome size

In order to examine the species-specific performance, we compared the prediction accuracy of EMPRES models with PhytoExpr B and C models within each species and across all CV folds. Within each species, EMPRES models using PlantCaduceus embeddings (EMPRES 1–3) consistently outperform the PhytoExpr benchmarks, with EMPRES 2 showing the highest accuracy in nearly all species (Figure 5b).

The magnitude of improvement by our top performing model (EMPRES 2) over the best benchmark (PhytoExpr C) varies across species. The smallest improvement is observed in *Medicago truncatula* (*Mtr*), where prediction accuracy increases from *R* = 0.78 to *R* = 0.84 (Δ*R* ≈ 0.06), corresponding to an increase in variance explained (Δ*R*^2^) of approximately 9.4%. In contrast, the largest improvement is observed in *Chlamydomonas reinhardtii* (*Cre*), where prediction accuracy increases from *R* = 0.55 to *R* = 0.76 (Δ*R* ≈ 0.21), corresponding to an increase in variance explained (Δ*R*^2^) of approximately 27%. Together, the results from sections 3.2 and 3.3 suggest that sequence embeddings from PlantCaduceus provide useful information for prediction of mRNA abundance. They also indicate that even though chromatin accessibility embeddings (from a2z) capture useful information, they are not as useful as PlantCaduceus gLM embeddings.

Finally, the comparison of species-specific prediction accuracy with genome size reveals that there is no direct relationship between genome size and the accuracy of gene expression prediction (Figure 5c). EMPRES 2 achieves the highest prediction accuracy in *Oryza sativa* (*Osa*) followed by *Zea mays* (*Zma*), despite large differences in genome size. Notably, EMPRES 2 has different prediction accuracy in species with similar genome size (*Arabidopsis thaliana* vs. *Chlamydomonas reinhardtii*, *Oryza sativa* vs. *Populus trichocarpa*, etc.). This suggests that some other factors, e.g., the quality of whole genome sequencing (WGS), gene annotations, and the percentage of repetitive sequence (such as transposons) of genome might be more related to prediction accuracy than genome size.

### 3.4 EMPRES models can accurately predict between-gene expression differences among control lines in a Brachypodium mutant population

In order to examine the accuracy of our proposed models in an unseen population, the models were used to generate predictions for expression levels of 23,725 genes in the control lines of the SIEVE population (Figure 3). Predictions of top five models for each four types of our EMPRES models were averaged in each CV fold (ensembling), and our ensembled predictions were compared with the observed expression averaged over control lines. EMPRES 1 and EMPRES 2 achieve the highest prediction accuracy, as quantified by the regression coefficient estimate *β*_1,*m*_ from equation (1) among all models included in the comparison (Table 2).

**Table 2:** Experimental validation results from between-gene and within-gene expression predictions on SIEVE Bdi population using our proposed EMPRES models and benchmark models. The prediction accuracy is quantified by the regression coefficient *β*_1,*m*_ for between-gene and within-gene predictions from equations (1) and (3), respectively.

| Source of variation | S2E model | Prediction accuracy | Standard error | P-value | R<sup>2</sup> |
| --- | --- | --- | --- | --- | --- |
| **Between Genes**<br>(Control lines) | PhytoExpr B | 0.556 | 0.0066 | $<2.2 \times 10^{-16}$ | 0.228 |
| | PhytoExpr C | 0.569 | 0.0066 | $<2.2 \times 10^{-16}$ | 0.239 |
| | EMPRES 1 | 0.778 | 0.0064 | $<2.2 \times 10^{-16}$ | 0.381 |
| | EMPRES 2 | 0.777 | 0.0064 | $<2.2 \times 10^{-16}$ | 0.381 |
| | EMPRES 3 | 0.764 | 0.0068 | $<2.2 \times 10^{-16}$ | 0.349 |
| | EMPRES 4 | 0.671 | 0.0089 | $<2.2 \times 10^{-16}$ | 0.193 |
| **Within Genes**<br>(Mutant lines) | PhytoExpr B | 0.083 | 0.030 | $5.7 \times 10^{-3}$ | $1.4 \times 10^{-5}$ |
| | PhytoExpr C | 0.060 | 0.027 | $2.8 \times 10^{-2}$ | $9.0 \times 10^{-6}$ |
| | EMPRES 1 | 0.341 | 0.047 | $2.1 \times 10^{-13}$ | $1.0 \times 10^{-4}$ |
| | EMPRES 2 | 0.384 | 0.046 | $8.7 \times 10^{-17}$ | $1.3 \times 10^{-4}$ |
| | EMPRES 3 | 0.341 | 0.047 | $3.9 \times 10^{-13}$ | $9.8 \times 10^{-5}$ |
| | EMPRES 4 | 0.033 | 0.039 | 0.393 | $1.4 \times 10^{-6}$ |

Noticeably, all EMPRES models using PlantCaduceus embeddings (EMPRES 1-3) outperform the benchmark models. This superior performance confirms the advantage of using regulatory sequence embeddings generated by the PlantCaduceus gLM as features for inferring gene expression from sequence. This experiment on control lines of the SIEVE population serves as an independent validation of generalizability of EMPRES models to new populations for predicting between-gene expression.

### 3.5 EMPRES models can predict allelic differences in expression levels caused by point mutations in independent mutant lines

To validate the ability of our models to predict variant effect as single-base resolution, each model was first used to predict the difference in expression for a given gene between mutant and control lines (within-gene variation). Each model’s predicted deviation (the difference between the prediction in mutant lines and the average prediction among control lines) was then compared to the observed deviation for each gene, using multiple linear regression models. Latent factors and potential confounders that could alter gene expression were accounted for in the regression using an optimal number of principal components as covariates.

The results show that predictions of EMPRES 1-3 models have a highly significant and positive association with observed expression changes, with EMPRES 2 model predictions being the most significant among all models (Table 2). The positive coefficients of EMPRES 1-3 models confirm that an increase in the model’s predicted expression is associated with an increase in observed gene expression in the mutants. Our models outperformed the benchmarks on predicting allelic differences in RNA abundance in the SIEVE population (regression coefficient β=0.38 for EMPRES 2 vs. β=0.06 for PhytoExpr C), despite being trained on the same data. Nonetheless, the accuracy gap between allele-level and gene-level predictions (e.g., for EMPRES 2: β=0.38 for allele-level vs. β=0.78 for gene-level) highlights significant room for improvement in variant effect prediction by S2E models.

Although EMPRES 1-3 models show statistically significant regression coefficients for within-gene expression changes, the proportion of variance explained (R^2^) remains low (Table 2). Importantly, 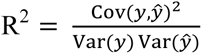 values are not directly comparable between within-gene and between-gene analyses, because the error variance of the response variable differs substantially between these two tasks (much more noise due to non-genetic and trans genetic variation in within-gene vs. between-gene expression). Moreover, predicted allelic deviations exhibited much lower variance than gene-level expression averages (Figure 3b). In this context, the regression coefficient *β*_1,*m*_ from equations (1) and (3) provides a more appropriate and comparable measure of predictive signals, as it quantifies the covariance between predicted and observed expression changes normalized by the variance of the predictions rather than the response. The significant and positive *β*_1,*m*_ estimates therefore indicate that the models capture a real, non-random genetic signal at the allelic level, despite the higher noise inherent to deviations of individual mutant lines from the control average.

## 4. Discussion

### 4.1 Feature contributions: Regulatory sequence embeddings and chromatin state

Overall, the EMPRES models leveraging PlantCaduceus sequence embeddings achieved the strongest performance across both cross-species evaluations and independent *in planta* validation. Among the models, EMPRES 2 showed the most consistent performance, indicating that PlantCaduceus embeddings encode regulatory information relevant to between-gene and within-gene expression differences, although predictive accuracy was reduced for allelic variation. Moreover, EMPRES 4, which relies solely on chromatin accessibility features, remained competitive with existing benchmarks, confirming that chromatin state alone is an informative predictor of gene expression [26], [39]. Together, these results indicate that both regulatory sequence embeddings and chromatin-related features contribute meaningful information, but that their naive combination may not be optimal, suggesting that more structured integration strategies could further improve predictive performance. A possible alternative approach could be to first train models on gene expression using gLM embeddings as features and then fine-tune the models by incorporating chromatin accessibility as additional features [39].

Our results differ from [40] on the representational power of pre-trained models in regulatory genomics. We showed that gLMs embeddings, without any fine-tuning of the gLM for downstream tasks, can be useful in regulatory genomics on a large-scale, cross-species plant dataset. The differences between our results and those of [40] may originate from the fact that while [40] focused on cell-type-specific predictions for two specific cell types (HepG2 and K562), we did not restrict our predictions and comparisons to any cell types. Other differences could stem from using a different gLM in our study vs gLMs in [40] with different length of input sequence (few hundreds of bps vs 10000 bps in our study) and prediction tasks, as we did not train or evaluate our models for tasks such as predicting enhancer activity and TF binding sites.

The lower predictive performance of EMPRES 3 in comparison with EMPRES 1 and EMPRES 2 for between-gene variation differs from the results reported by DeepCBA. However, DeepCBA does not use sequence embeddings from a sequence-to-chromatin model, as leveraged in EMPRES 3, but relies on explicitly defined chromatin interaction structure. Therefore, the two approaches employ fundamentally distinct forms of chromatin information, which may contribute to the different results observed across studies.

More broadly, improving S2E models often involves a trade-off between feature richness and model generalizability. While incorporating additional epigenomic or regulatory features can enhance predictive performance [24], [41], [42], such approaches increase computational and data requirements and typically rely on species-specific resources, limiting scalability across species [14]. In contrast, EMPRES prioritizes cross-species generalizability by relying on sequence-derived embeddings, a deliberate design choice that favors broad applicability over maximal, species-specific accuracy.

### 4.2 Model training specifically for variant effect prediction

The validation task in this study was challenging, especially for predicting allelic differences, where the models trained exclusively on reference genomes were used to predict the effect of single nucleotide mutations *in planta*—an extremely subtle signal within a 10 kb sequence input. Another challenge in our validation was using leaf tissue expression values in the SIEVE dataset, while our models were trained on median expression values of samples across all tissues, similarly to [1]. In this validation task, our proposed EMPRES models achieved high statistical significance (*p* << 0.05) in associating predicted expression changes with observed changes, demonstrating they capture a true genetic signal. On the other hand, there was a substantial decrease in the regression coefficient of all our proposed models from between-gene variation to within-gene variation (Table 2, Figure 6). Interestingly, for predicting variant effects, EMPRES 2 model had the highest regression coefficients among all models, and all EMPRES models using PlantCaduceus embeddings outperformed competitive models (Table 2).

**Figure 6:**
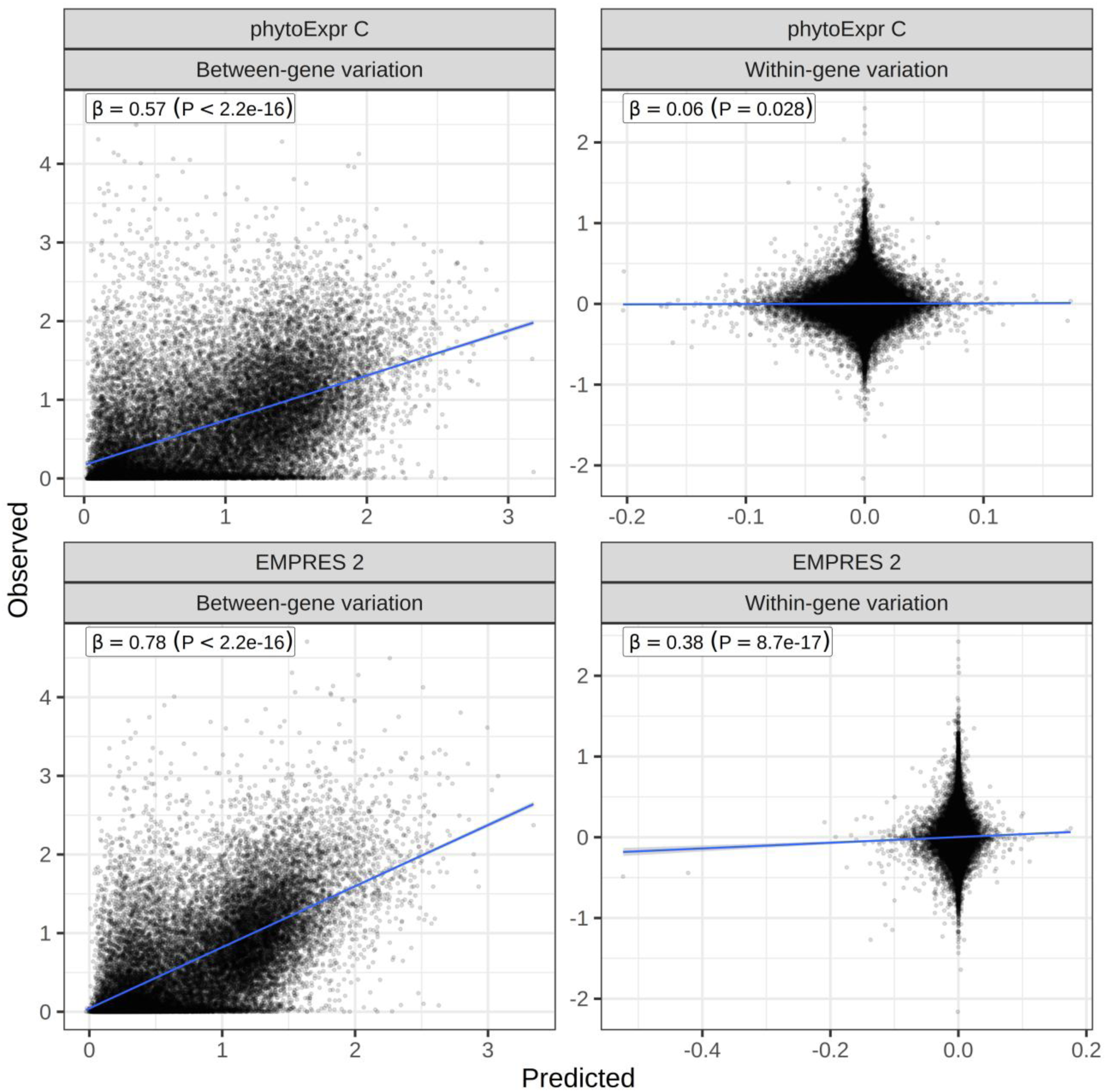
Validation of predicted mRNA abundance in the SIEVE population. Scatter plots show the relationship between predicted and observed mRNA abundance in *Brachypodium distachyon* SIEVE lines for PhytoExpr C (top row) and EMPRES 2 (bottom row). Left panels demonstrate between-gene variation, comparing predicted and observed average expression levels across control lines for each gene. Right panels demonstrate within-gene variation, comparing predicted and observed deviations in expression between mutant lines and the corresponding gene-specific control mean. Observed and predicted expression values are shown on the log_10_ (1+TPM) scale for between-gene analyses, and as expression deviations for within-gene analyses, as defined in Methods. Each point represents one gene (between-gene) or one gene–mutant observation (within-gene). Blue lines indicate fitted linear regression models. The regression coefficient β and associated P-value (denoted by P) are reported for each panel, corresponding to *β*1,*m* in equation (1) for between-gene and equation (3) for within-gene analyses.

Our approach is different from foundational models such as AlphaGenome [25], which is a massive-scale, unified and multi-task model, highly capable of zero-shot variant effect prediction across different modalities such as gene expression, chromatin accessibility, histone modification, etc., at single-base resolution. Our approach also differs from plant-specific sequence-to-expression models such as DeepWheat [43], which uses a two-stage architecture, where epigenomic features are first predicted from DNA sequence and then integrated with sequence information to predict gene expression and variant effects in a single crop species. Such approaches leverage rich, species-specific epigenomic datasets to guide expression prediction, but require direct supervision from epigenomic measurements during training.

EMPRES also differs from other novel S2E models such as DeepAllele [44], which are trained on allele-specific expression data to directly predict expression changes caused by allelic variations. Variant effect prediction studies such as [44], [45] use contrastive learning to fine-tune pretrained models on individual-specific paired genomic and transcriptomic data to achieve high predictive power for effects of mutations on gene expression. The ability of our proposed models to capture a true and statistically significant signal without such personalized data or modeling strategies is therefore a notable achievement and establishes a powerful baseline for future optimizing efforts. Future efforts could consist of combining sequence embeddings (e.g., from PlantCaduceus) with a contrastive learning approach and, if possible, allele-specific expression data. Unlike [1] which could employ gradient-based attribution methods to assign importance scores to individual nucleotides, our approach makes gradient-based attribution difficult or even unfeasible. However, perturbation-based attribution methods (*in silico* mutagenesis) were still possible with our approach (e.g., in the SIEVE validation). Moreover, more advanced *in silico* model explanation toolkits for cis-regulatory region such as CREME [46] may be applied to better understand how variations in input sequences affect model predictions.

### 4.3 Overcoming the computational cost of training on pre-trained gLM embeddings

A practical limitation of our pipeline is its computational burden, driven by large-scale feature generation with pretrained gLMs, and model training. Given the strong performance in this study achieved by EMPRES models using PlantCaduceus embeddings, a possible direction for improving scalability is knowledge distillation (KD). In KD, a compact “student” model is trained to approximate a high-performing “teacher” model’s outputs rather than the labels directly [47]. KD has been used in regulatory genomics together with ensemble learning to compress accurate predictors while reducing inference cost [48]. In our setting, the teacher could be an EMPRES 2 ensemble, and the student could use a cheaper input representation. Then, the student–teacher divergence can be quantified directly as the prediction loss on held-out genes (e.g., MSE between student and teacher predicted expression). Key challenges include the cost of generating teacher predictions at scale and the extent to which a student trained on simpler representations can reproduce the teacher’s behavior while retaining predictive performance. Addressing these challenges will be an important direction for future work aimed at improving the scalability and practical applicability of S2E models.

### 4.4 Conclusions: EMPRES is an effective approach for predicting gene expression in plants

This study demonstrates that leveraging pre-trained gLMs can provide a substantial leap forward in predicting gene expression from proximal regulatory sequences in plants. Our proposed EMPRES models, using the PlantCaduceus gLM embeddings, significantly outperform the benchmark models based on one-hot encoding, for predicting between-gene expression across 17 plant species. Moreover, in a challenging *in planta* validation, our models successfully predicted the directional effect of single point mutations on gene expression with high statistical significance, a task where the benchmark models failed, thereby addressing a critical gap in the field. Overall, this work introduces and validates a new paradigm for S2E modeling in plants, shifting from direct sequence encoding to the use of information-rich, pre-trained gLM embeddings.

Beyond improved predictive performance, our work highlights the value of carefully designed experimental resources, such as the SIEVE mutant population, for discriminating models based on their ability to capture allelic effects at single-base resolution in a controlled genomic background. Our approach shows opportunities for plant biologists and the machine learning community to predict regulatory variant effects *in planta* for potential applications in precision breeding and crop improvement. It points to one important direction for future progress—using informed sequence embeddings rather than naive sequence encoding—among others (e.g., additional data modalities and alternative modeling strategies), which together could help close the gap between allele-level and gene-level prediction accuracy.

## Acknowledgement

This research is supported by the Novo Nordisk Foundation, Grant NNF21OC0067311. We would like to thank GenomeDK and Aarhus University for providing computational resources and support that contributed to these research results.

## Data availability statement

All data used in this study, including sequence embeddings, trained models, predicted expression values, and supplementary analysis files, are publicly available via Zenodo at https://doi.org/10.5281/zenodo.18236856. All scripts used to run the analyses and train the supervised models are available at https://github.com/behroozvahedi/Regeffects GitHub repository.

## Abbreviations

CRE: cis-regulatory element
TSS: transcription start site
TTS: transcription termination site
TF: transcription factor
CNN: convolutional neural networks
TPM: transcript per million
gLM: genomic language model
S2E: sequence-to-expression
eQTLs: expression quantitative trait loci
CV: cross-validation
PC: principal component
BIC: Bayesian information criterion
WGS: whole genome sequencing
KD: knowledge distillation

