## Supplementary material for "Genomic language models improve cross-species gene expression prediction and accurately capture regulatory variant effects in Brachypodium mutant lines"

**Table S1:** Space of prior values used by Optuna's TPE sampler to select hyperparameter values for minimizing the validation loss.

| **Hyperparameter** | **Range of Values** | **Type** |
|---|---|---|
| convolutional layers | {3, 4, 5} | categorical |
| Kernels (filters) | {128, 192, 256, 320, 384} | categorical |
| Kernel size | {1, 2, 3, 4} | categorical |
| Dense layers | {3, 4, 5} | categorical |
| Dense layers after merging branches | {2, 3, 4} | categorical |
| Dense layers size | {16, 32, 64, 128} | categorical |
| Batch size | {64, 128, 256} | categorical |
| Dropout rate | [0.0, 0.5] | continuous |
| Learning rate | [1e-5, 1e-2] | continuous |

a)

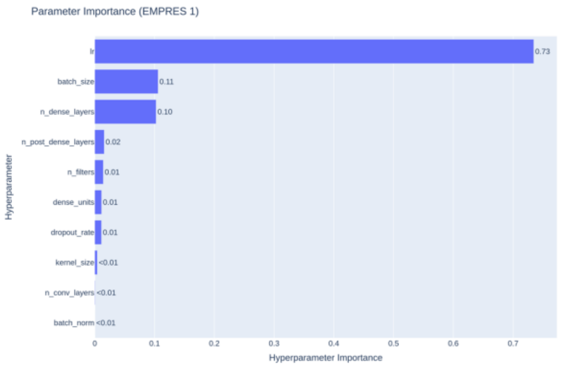


b)

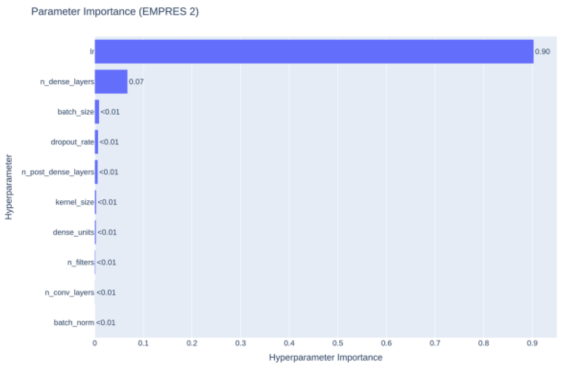


c)

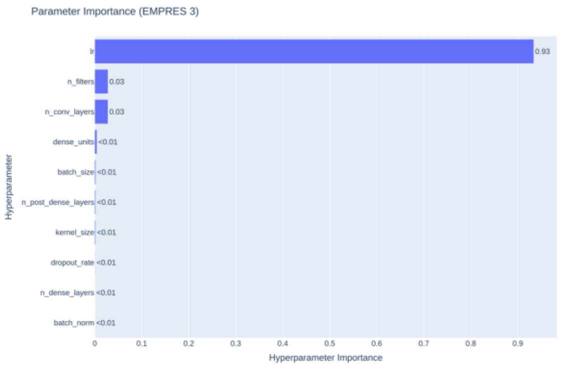


d)

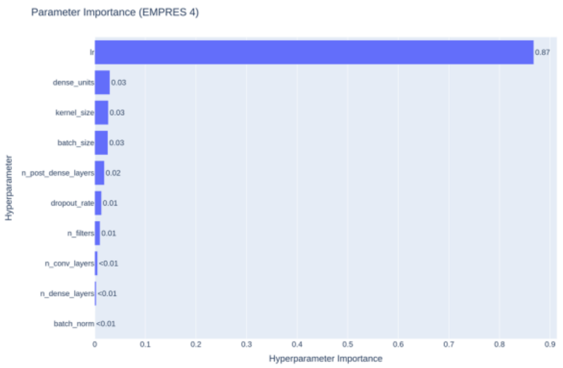


**Figure S1:** Optuna parameter importance plot for each EMPRES type, showing the relative importance of hyperparameters for optimizing the performance of models during training

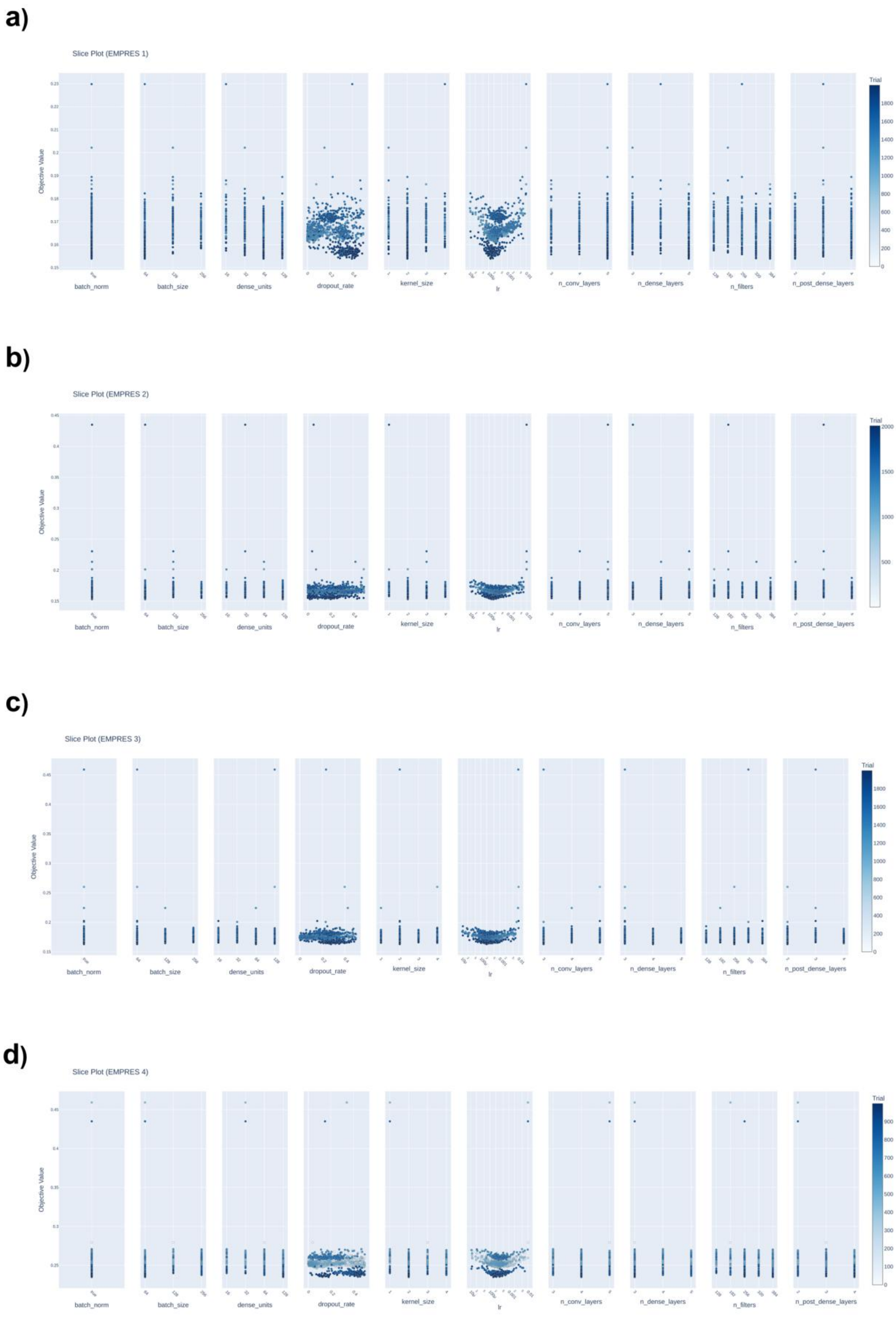


**Figure S2:** Optuna slice plot showing the distribution of hyperparameter values and the corresponding validation MSE loss for each model. a) EMPRES 1, b) EMPRES 2, c) EMPRES 3, d) EMPRES 4

**a)**

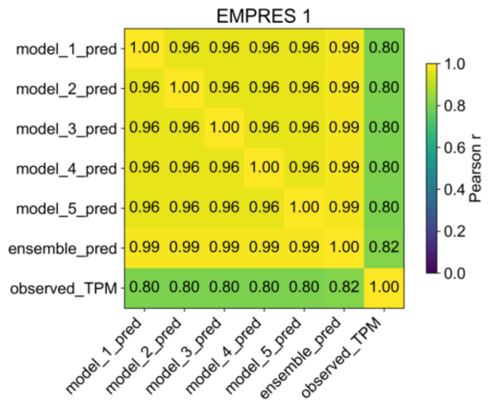


**b)**

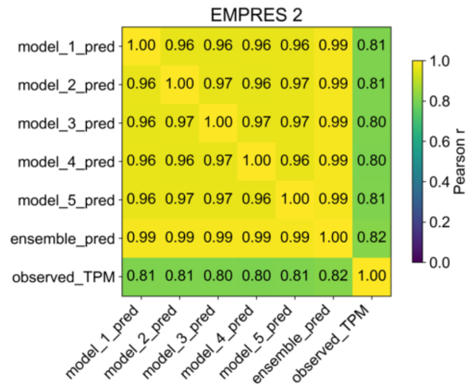


**c)**

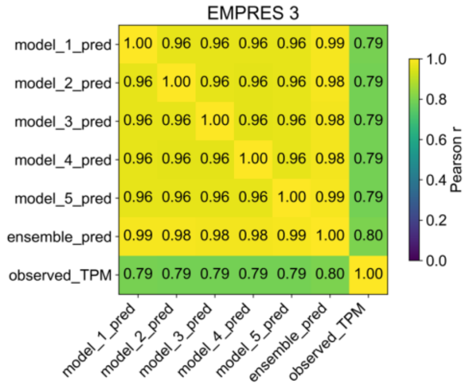


**d)**

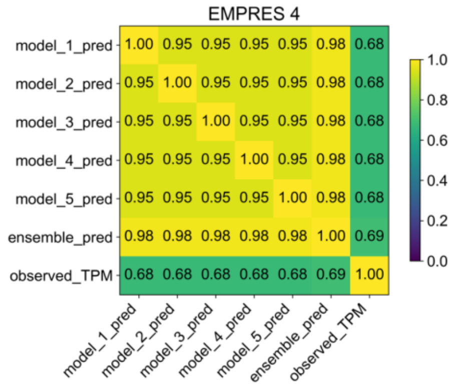


**Figure S3:** Heatmaps of prediction accuracy defined as the Pearson correlation coefficient between predicted and observed TPM value, for the top five models and the ensemble of top 5 models to investigate the effect of ensembling.

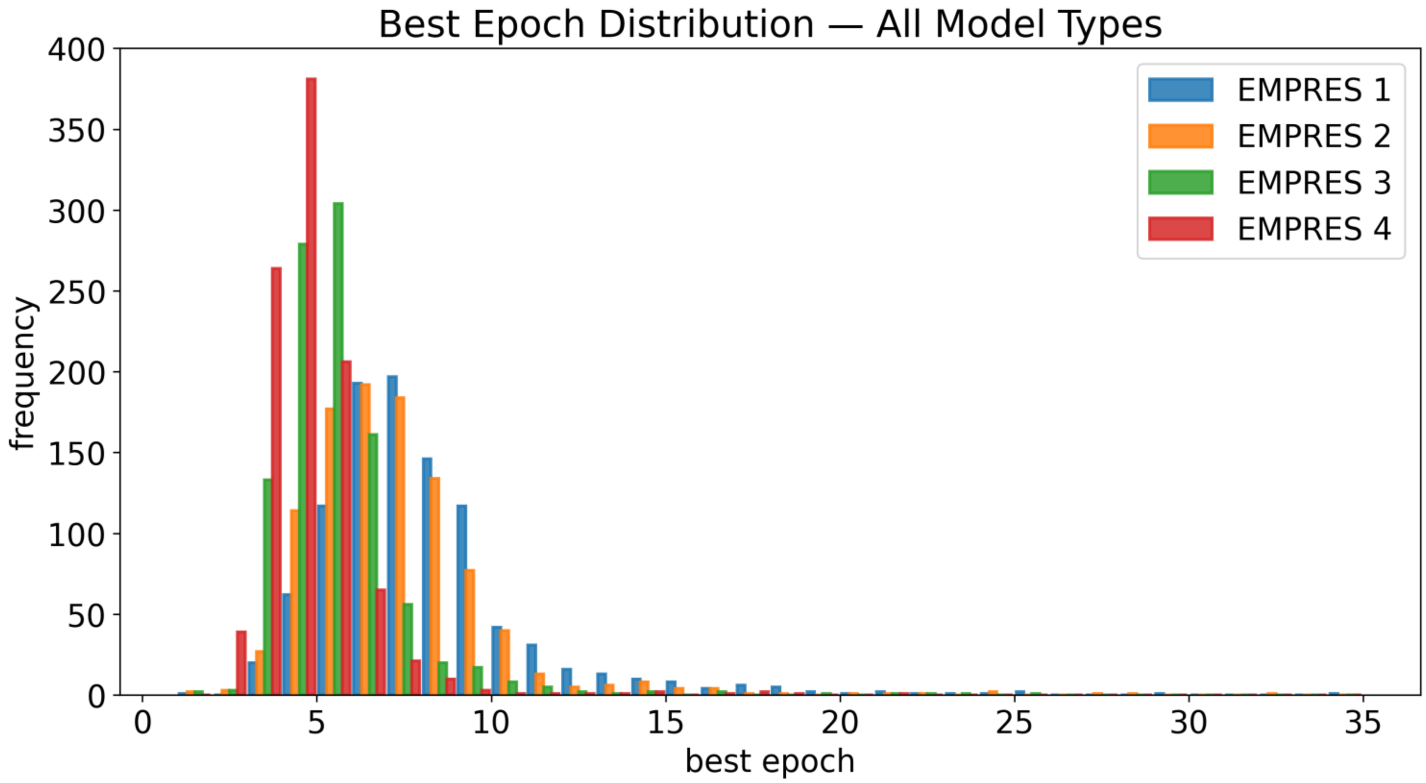


**Figure S4:** Distribution of best epoch for all trained models across all CV folds for each EMPRES type. For each model, the best epoch is determined as the epoch with minimum MSE loss between predicted and observed gene expression in the validation dataset for each CV fold.
